# Minimal models of the inspiratory and sigh breathing rhythms of the preBötzinger complex

**DOI:** 10.1101/2022.11.15.516637

**Authors:** Daniel S. Borrus, Cameron J. Grover, Christopher A. Del Negro, Gregory D. Conradi Smith

**Author notes:** Corresponding authors contributed equally: Gregory D. Conradi Smith, Ph.D., Christopher A. Del Negro, Ph.D.

## Abstract

The preBötzinger complex of the lower brainstem generates two breathing-related rhythms: one for inspiration on a timescale of seconds, and another that produces larger amplitude sighs on the order of minutes. We hypothesize that these two disparate rhythms emerge in tandem wherein recurrent excitation gives rise to the inspiratory rhythm while a calcium oscillator generates sighs; distinct neuronal populations are not required. We present several mathematical models that instantiate our working hypothesis including: (1) an activity (firing rate) model and (2) a minimal spiking network model. Both modeling frameworks corroborate the single-population rhythmogenic hypothesis.

## 1 Introduction

The preBötzinger complex (preBötC) of the lower brainstem generates two breathing-related rhythms: one for inspiration on a timescale of seconds, and another that produces larger amplitude sighs on the order of minutes. Their underlying mechanisms and cellular origins remain incompletely understood.

This report summarizes the development of several mathematical models designed to test the consistency of the hypothesis that the inspiratory and sigh rhythms may emerge from a single population of neurons. In this view, recurrent excitation gives rise to the inspiratory rhythm while intracellular calcium oscillations generate sighs. Using an activity (firing rate) model and, subsequently, a minimal spiking network model, simulation results corroborate the hypothesis that both the inspiratory and sigh rhythms may emerge from a single population of excitatory preBötC interneurons.

The remainder of this document is organized as follows. Section 3 focuses on the activity model of episodic inspiratory burst generation. Section 4 presents the dynamical model of slow Ca^2+^ oscillations that drive the sigh rhythm. Section 5 gives details regarding the coupling and interaction of these two subsystems. Section 6 compares the activity model of inspiratory rhythmogenic preBötzinger complex neurons with a spiking network model that includes Ca^2+^ dynamics for several hundred distinct neurons with excitatory interactions mediated by a physiologically plausible network topology.

## 2 Description of model of the inspiratory and sigh rhythms

The activity model of inspiratory and sigh rhythmogenesis (Fig. 1) is an ordinary differential equation (ODE) system with five dynamical variables—three for the inspiratory activity of the preBötzinger complex (preBötC) neuronal network (*a, s, θ*) and two for the dynamics of intracellular calcium (Ca^2+^) in a representative preBötC neuron (*c, c*_*tot*_). The differential equations for the full model are

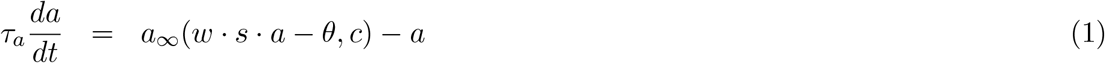

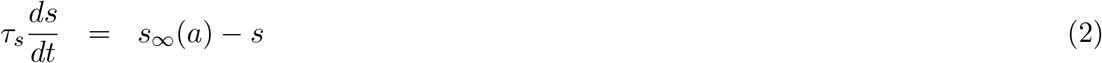

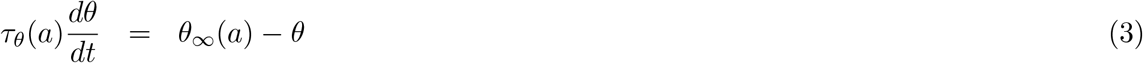

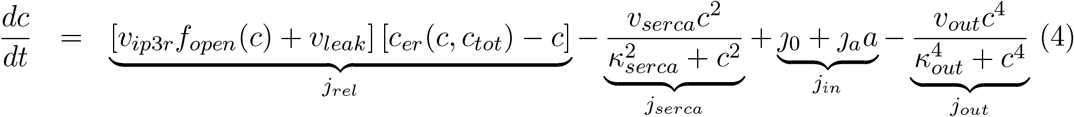

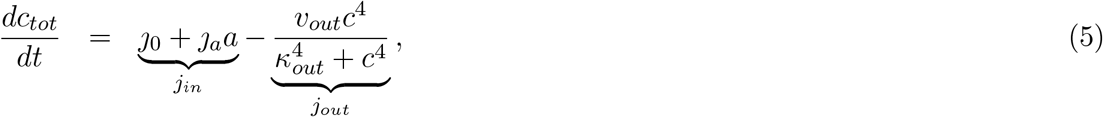

where *j*_*rel*_ and *j*_*serca*_ are endoplasmic reticulum Ca^2+^ fluxes, and *j*_*in*_ and *j*_*out*_ are plasma membrane Ca^2+^ fluxes. The functions *a*_∞_, *s*_∞_, *θ*_∞_ and *τ*_*θ*_ that appear in the epnea subsystem (Eqs. 1–3) are

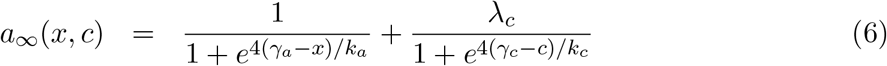

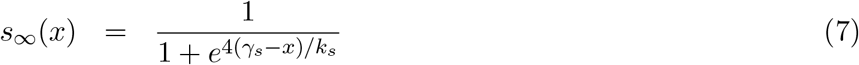

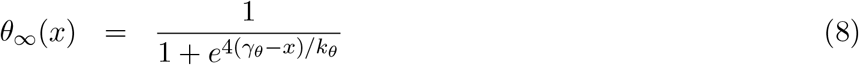

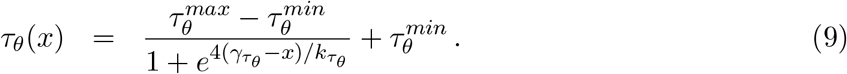

**Figure 1:**
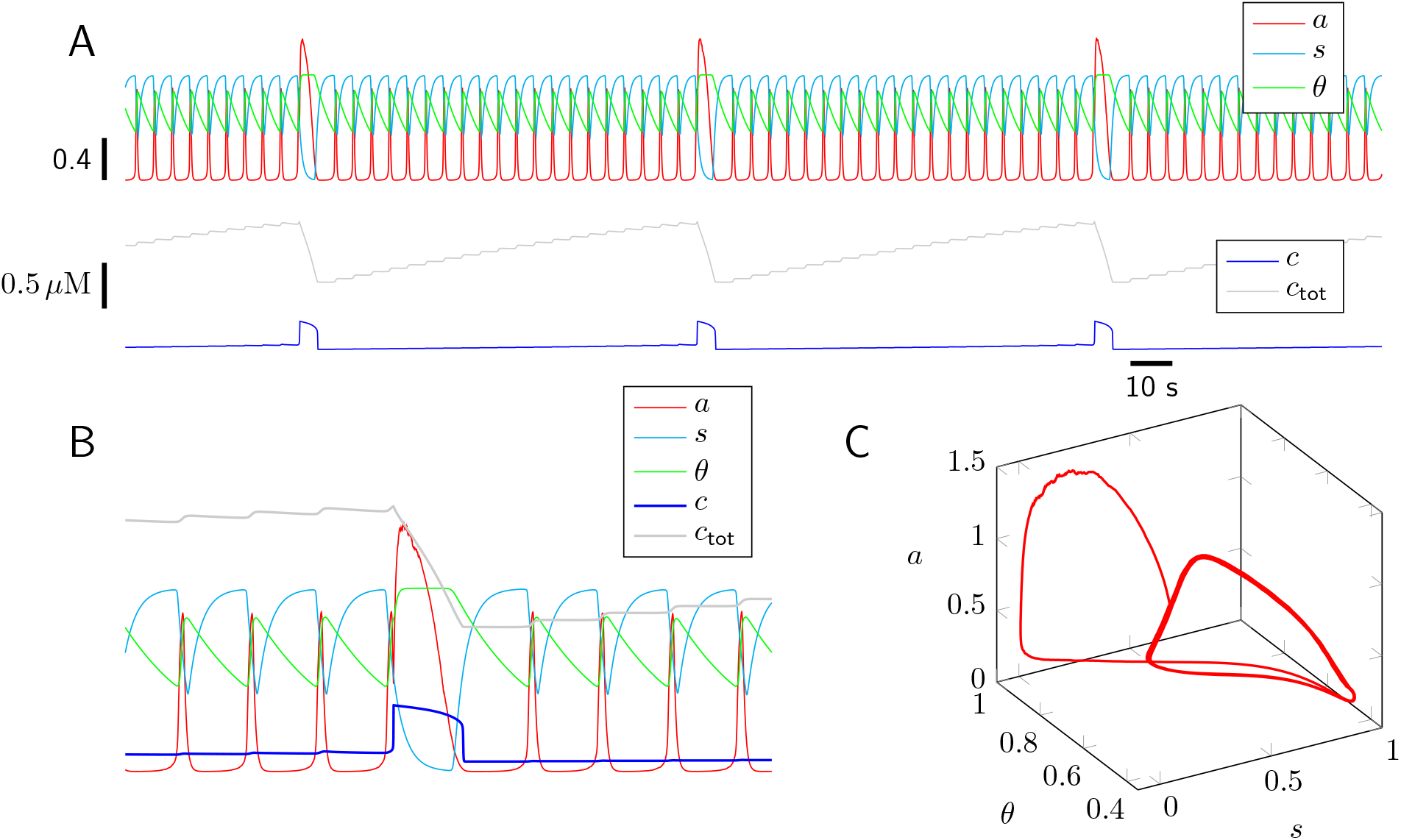
Representative simulation of the coupled inspiratory and sigh rhythms. (A) Dynamics of the five dependent variables of the ODE model. (B) Expanded view that includes a single sigh event that emerges shortly after the onset of inspiratory burst. (C) Three dimensional phase space for the (*a, s, θ*) subsystem. Parameters as in Tables 1–2.

The first state variable, denoted by *a*, is the network activity of the preBötC. This dimensionless quantity takes values between 0 (no activity) and 1 + *λ*_*c*_ (maximum population firing rate). The dimensionless variables, *s* and *θ*, model the dynamics of synaptic depression and cellular adaptation, respectively. The variable *c* is the cytosolic free Ca^2+^ concentration (*c* = [Ca^2+^]). The variable *c*_*tot*_ is the total [Ca^2+^], a quantity dominated by intracellular stores. Denoting endoplasmic reticulum (ER) [Ca^2+^] as *c*_*er*_, the total [Ca^2+^] is defined as *c*_*tot*_ = *c* + *ρc*_*er*_, where where *ρ* is an ER-to-cytosol volume ratio that accounts for the Ca^2+^-buffering capacity of both compartments. There are two algebraic functions that occur in Eq. 4. The first expresses the ER [Ca^2+^] in terms of the cytosolic and total [Ca^2+^],

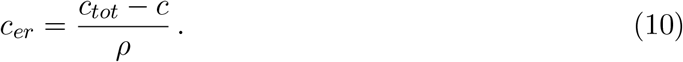

The second function is the bell-shaped equilibrium open probability of the IP_3_ receptor (IP_3_R) as a function of cytosolic [Ca^2+^],

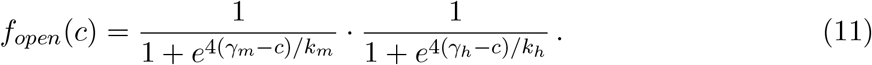

See Tables 1–2 for a description of parameters and their standard values.

**Table 1:**
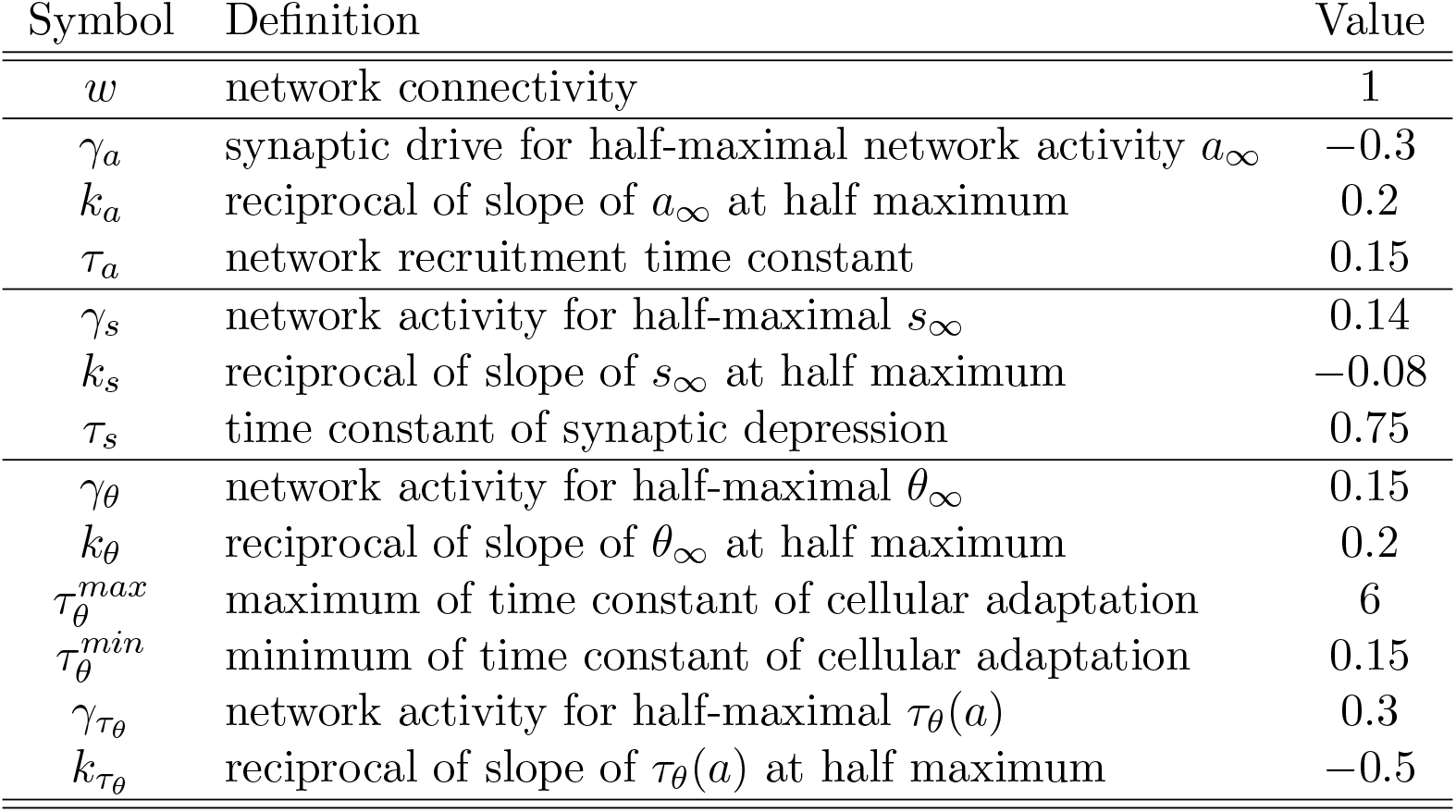
Standard parameters for inspiratory model (dimensionless).

**Table 2:** Standard parameters for Ca^2+^ subsystem.

| Symbol | Definition | Value | Units |
| --- | --- | --- | --- |
| $v_{ip3r}$ | rate constant of $\text{Ca}^{2+}$ release | 20 | $\text{s}^{-1}$ |
| $v_{leak}$ | rate constant of $\text{Ca}^{2+}$ leak | 0.25 | $\text{s}^{-1}$ |
| $v_{serca}$ | maximum rate of SERCA pumps | 60 | $\mu\text{M s}^{-1}$ |
| $\kappa_{serca}$ | half maximum for SERCA pumps | 0.3 | $\mu\text{M}$ |
| $\rho$ | ER/cytosol effective volume ratio | 0.15 | - |
| $\gamma_m$ | activation of intracellular $\text{Ca}^{2+}$ channels | 0.25 | $\mu\text{M}$ |
| $k_m$ | reciprocal of slope of $m_\infty$ at half maximum | 0.16 | $\mu\text{M}^{-1}$ |
| $\gamma_h$ | activation of intracellular $\text{Ca}^{2+}$ channels | 0.3 | $\mu\text{M}$ |
| $k_h$ | reciprocal of slope of $h_\infty$ at half maximum | -0.24 | $\mu\text{M}^{-1}$ |
| $j_0$ | constant $\text{Ca}^{2+}$ influx rate | 0 | $\mu\text{M s}^{-1}$ |
| $j_a$ | $\text{Ca}^{2+}$ influx rate proportionality constant | 0.1 | $\mu\text{M s}^{-1}$ |
| $v_{out}$ | maximum rate of PMCA pumps | 0.4 | $\mu\text{M s}^{-1}$ |
| $\kappa_{out}$ | half maximum for PMCA pumps | 0.3 | $\mu\text{M}$ |
| $\lambda_c$ | maximum $\text{Ca}^{2+}$-dependent increase of activity | 1.5 | - |
| $\gamma_c$ | threshold for $\text{Ca}^{2+}$-dependent increase of activity | 0.33 | $\mu\text{M}$ |
| $k_c$ | reciprocal slope of $\text{Ca}^{2+}$-dependent increase of activity | 0.05 | $\mu\text{M}$ |

## 3 Activity model of inspiratory rhythm

Episodic bursting of the preBötC is modeled as a three-variable dynamical system,

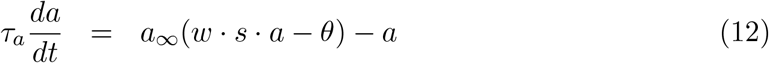

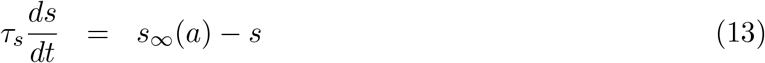

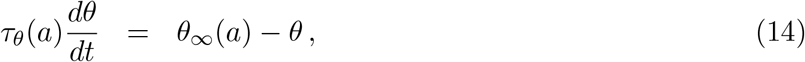

where the state variable *a* is the network activity, *s* accounts for the dynamics of synaptic depression (*s* = 1 indicates the absence of depression, while *s* = 0 corresponds to full depression), and *θ* represents activity-dependent cellular adaptation. The functions *a*_∞_ and *s*_∞_ in Eqs. 12 and 13 are sigmoidal,

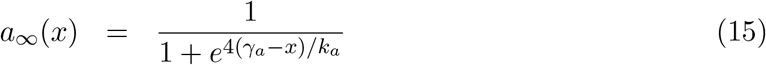

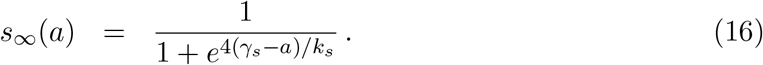

Because *k*_*a*_ is positive, *a*_∞_ is a monotone increasing function of *x* (Eq. 15), which in Eq. 12 is given by *x* = *w* · *s* · *a* − *θ*. In this expression, *w* is the synaptic gain, the triple product *w* · *s* · *a* is the aggregate synaptic drive, which is offset by cellular adaptation, *θ*). Conversely, *k*_*s*_ is negative; hence, *s*_∞_ in Eq. 13 is monotone decreasing function of the network activity (*a*). The steady-state level of cellular adaptation,

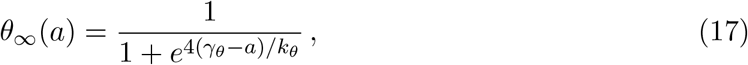

is an increasing function of network activity (*k*_*θ*_ > 0). The time constant for cellular adaptation, *τ*_θ_, is also a function of network activity,

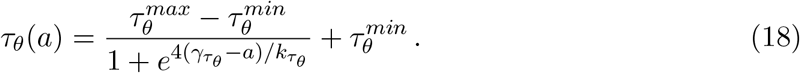

The 4 that appears in Eqs. 15 and 17 makes *k*_*a*_, *k*_*s*_ and *k*_*θ*_ the inverse of the slope of *a*_∞_, *s*_∞_, *θ*_∞_ at their respective half-maxima.

For a fixed amount of cellular adaptation (*θ* = *θ*_0_), steady state network activity (*ā*) solves

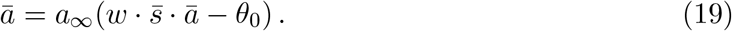

Fig. 2 shows that network activity can have 1–3 steady states (solutions of Eq. 19) depending on the amount of synaptic depression (*s*). For intermediate values of *s* (middle panel), the excitatory network is bistable. Fig. 3B shows how the increasing cellular adaptation has the effect of shifting the steady-state network activities to the right. That is, an increase in cellular adaptation (larger *θ*) can be offset by a decrease in synaptic depression (larger *s*). Parameters for the activity model of inspiratory rhythm (Table 1) are chosen to ensure that cellular adaptation accumulates rapidly in the active phase (large *a*), but recovers slowly during the silent phase when network activity is low, i.e., *τ*_*θ*_ is a decreasing function of *a* (Fig. 3A and 4C).

**Figure 2:**
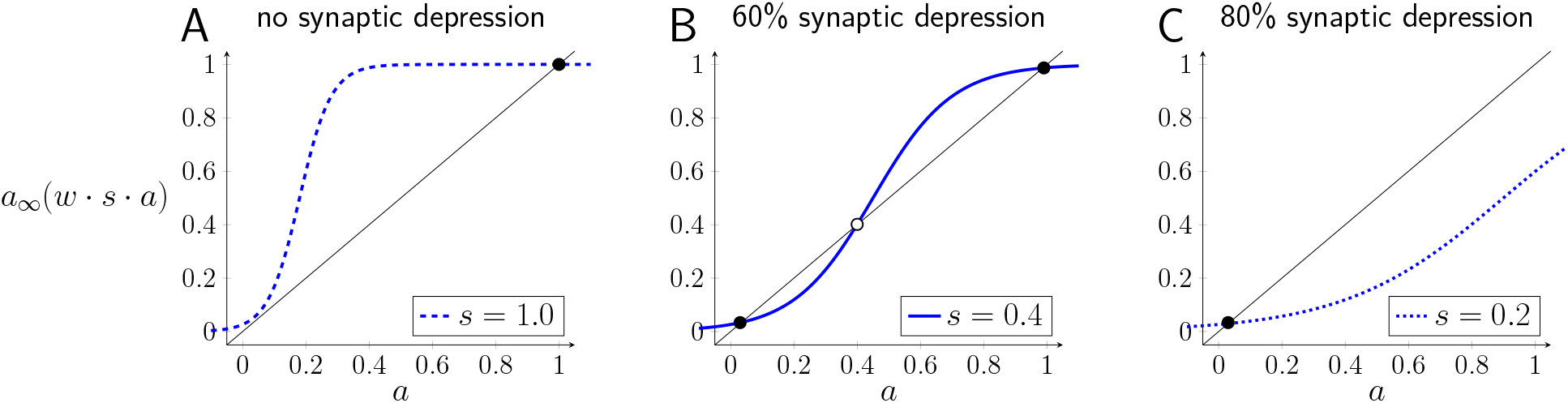
The recurrent excitatory network has steady-state population firing rates given by solutions of Eq. 19. The number of steady states depends on synaptic depression (*s*), which takes values between 0 (complete depression) and 1 (no depression). Stable and unstable steady states are shown by filled and open circles, respectively. (A) In the absence of synaptic depression there is a steady state near the maximal population firing rate (*a* ≈ 1) at the intersection of *a*_∞_ (solid blue curve) and the 45^°^ line (black) for *a* = *a*_∞_(*wsa*). (B) For intermediate synaptic depression the network is bistable. (C) For highly depressed synapses there is a stable steady state at low population firing rate (*a* ≈ 0). Parameters: *w* = 1, *γ*_*a*_ = 0.18, *k*_*a*_ = 0.2.

**Figure 3:**
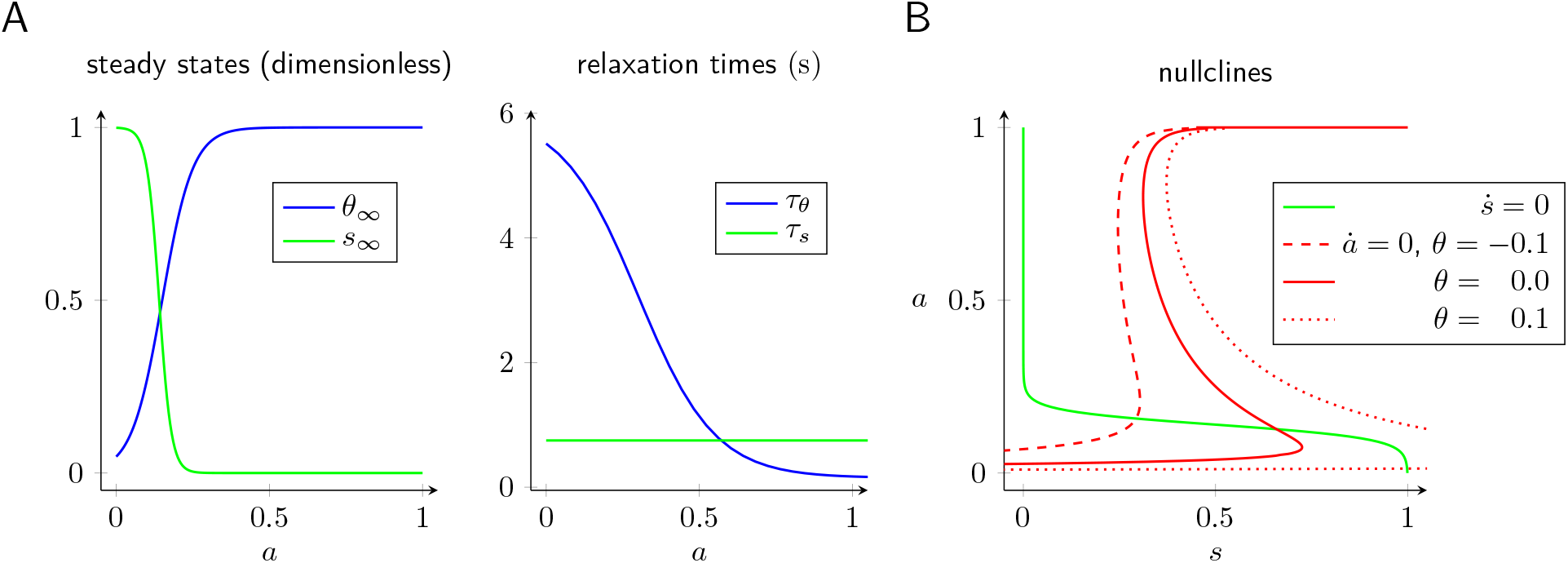
(A) Dynamics of synaptic depression (*s*) and cellular adaption (*θ*) depend upon network activity (*a*) via the steady-state functions, *s*_∞_ and *θ*_∞_, and exponential relaxation times, *τ*_*s*_ and *τ*_*θ*_. (B) Steady-state network activity for three different values of the variable representing cellular adaptation (*θ*, see legend).

Fig. 4A shows a representative simulation of the activity model of inspiratory rhythm. Synaptic depression terminates the bursts, but burst onset is determined by the recovery of cellular adaptation (*θ* must be sufficiently small), consistent with empirical data (Baertsch et al., 2018; Guerrier et al., 2015; Kottick and Del Negro, 2015). In particular, (1) the interburst interval distribution is normally distributed with realistic mean and variance (*µ* = 4.3 s, *s* = 0.044 s), and (2) there is no correlation between burst amplitude and the duration of the preceding inter-burst interval (*r*^2^ < 0.01) (cf. Fig. 4D and E).

**Figure 4:**
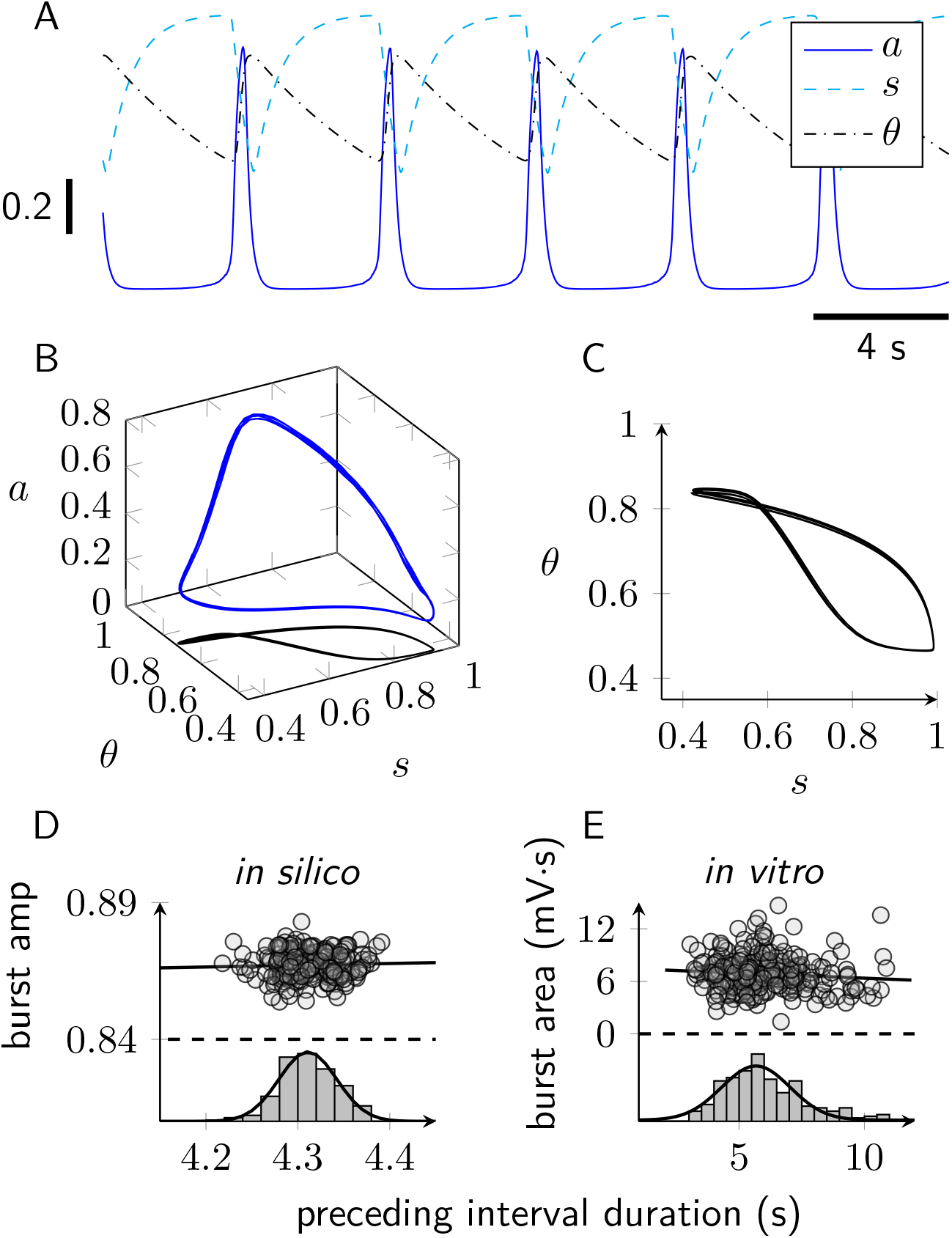
Three-variable model of episodic network activity gives realistic inspiratory dynamics (see Eqs. 12–14). (A) Representative trajectory of the three-state model showing network activity (*a*) and the dynamics of two slow variables: synaptic depression (*s*) and cellular adaptation (*θ*). (B) Trajectory in 3d phase space (blue curve, left) with projection emphasizing the relationship between synaptic depression and cellular adaption (*s* and *θ*, black curve, right). (C) Comparison of burst amplitude and inter-burst interval duration according to the model (in silico) and experiment (in vitro).

## 4 Ca^2+^ handling and the sigh rhythm

Fig. 1 shows a representative simulation of the coupled inspiratory and sigh rhythms with three sigh events. These occur because the inspiratory subsystem (*a, s, θ*) is coupled to oscillatory dynamics for intracellular Ca^2+^ (*c, c*_*tot*_) that periodically evoke a Ca^2+^-dependent increase in network activity (Eq. 6). The Ca^2+^ subsystem is the following two ODEs,

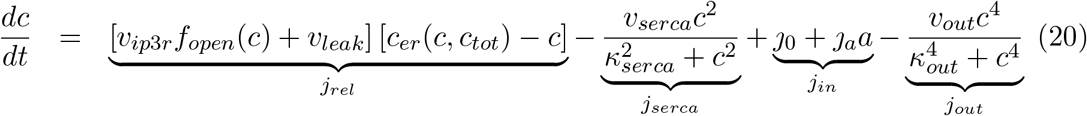

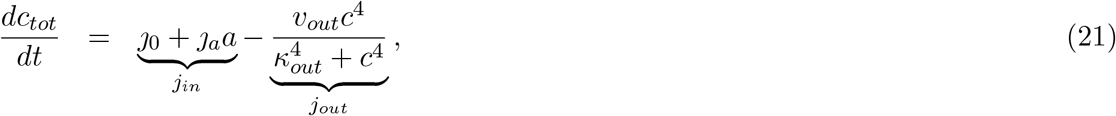

where the variable *c* is the cytosolic [Ca^2+^], and *c*_*tot*_ is the total intracellular [Ca^2+^] that includes contributions from the cytosol but is dominated by the [Ca^2+^] within the endoplasmic reticulum (ER). See (Friel and Chiel, 2008; Keizer et al., 1995) for review of Ca^2+^ dynamics modeled in this fashion.

The parameters *v*_*ip*3*r*_ and *v*_*leak*_ in Eq. 20 are rate constants for Ca^2+^-induced Ca^2+^ release and a passive leak, both with a driving force given by the concentration gradient across the ER membrane (*c*_*er*_ *− c*). The parameter *v*_*serca*_ is the maximal activity of a sarco-endoplasmic reticulum Ca^2+^ ATPase (SERCA) reuptake flux given by a sigmoidal Hill function with dissociation constant *κ*_*serca*_. The parameter *v*_*out*_ is the maximal activity of the sigmoidal expression *j*_*out*_(*c*) representing extrusion of Ca^2+^ by plasma membrane Ca^2+^ ATPases (PMCA). The terms *J* _0_ + *J*_*a*_*a* in Eqs. 20–21 model Ca^2+^ influx as a linear function of the network activity *a* (*J* _*a*_ is a propotionality constant and *J*_0_ is the background Ca^2+^ influx rate). The algebraic functions, *c*_*er*_(*c*) and *f*_*open*_(*c*), that occur in Eqs. 20 and 21) are given by *c*_*er*_ = (*c*_*tot*_ *− c*)*/ρ* and Eq. 11.

### Bistability in a closed cell model of Ca^2+^ handling

The relaxation oscillator dynamics of the Ca^2+^ subsystem that drive the sigh rhythm can be understood by considering how the dynamics of cytosolic [Ca^2+^] depends on total cell Ca^2+^ (*c*_*tot*_), which is the slow variable in Eqs. 20–21. If this slow variable were constant, the following ODE for the closed cell model would apply,

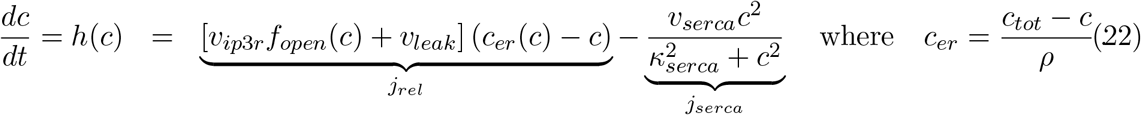

Using three different values for *c*_*tot*_, Fig. 5 plots the net ER flux *h*(*c*) = *j*_*rel*_ − *j*_*serca*_. For intermediate values of *c*_*tot*_ (solid blue curve), *h*(*c*) intersects the horizontal axis three times. Noting the slope *h*′(*c*) evaluated at these three steady states, it is evident that the low and high steady states are stable while the intermediate steady state is unstable.

**Figure 5:**
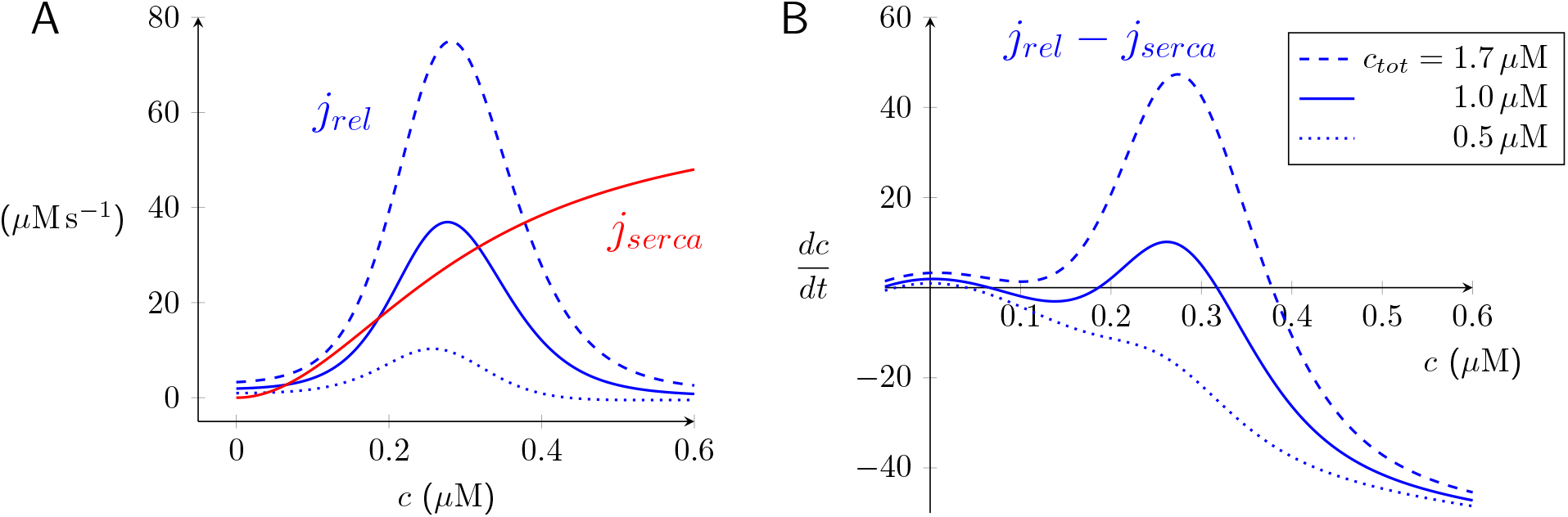
In the closed cell model of Ca^2+^ handling (Eq. 22) there are no plasma membrane fluxes; consequently, the total [Ca^2+^] (*c*_*tot*_) is parameter (not a state variable). (A) Fluxes associated to IP_3_R-mediated Ca^2+^ release (*j*_*rel*_) and SERCA pumps (*j*_*serca*_) as a function of cytosolic Ca^2+^ concentration (*c*). The release flux *j*_*rel*_ is an increasing function of *c*_*tot*_ because it is proportional to the concentration gradient (*c*_*er*_ − *c*) where *c*_*er*_ = (*c*_*tot*_ − *c*)/*ρ* (compare dotted, dashed, and solid curves; *c*_*tot*_ as in legend in B). The reuptake flux *j*_*serca*_ (shown red) is a function of *c* but not *c*_*tot*_. (B) Phase diagram of closed cell model of Ca^2+^ handling (Eq. 22). Because plasma membrane fluxes are not included in the close cell model, *c*_*tot*_ is constant (legend shows three values). Parameters as in Table 2.

### Relaxation oscillations in open cell model of Ca^2+^ handling

In the full model of coupled inspiratory and sigh rhythms (Eqs. 1–11), the fluxes representing Ca^2+^ release and reuptake from the ER (Eq. 22) are augmented by plasma membrane fluxes, *j*_*pm*_ = *j*_*in*_ *− j*_*out*_ where *j*_*in*_ = *j*_0_ + *J*_*a*_*a* and 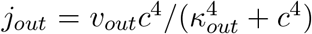. By averaging the Ca^2+^ influx over cycles of episodic bursting with period *T*, one may calculate an effective Ca^2+^ influx rate that no longer depends on the network activity (*a*),

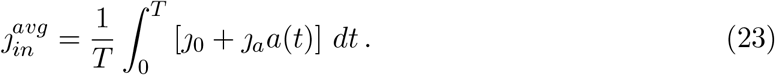

The resulting open cell model is

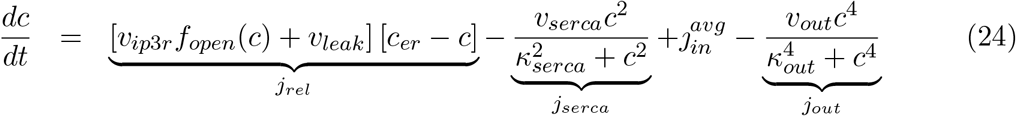

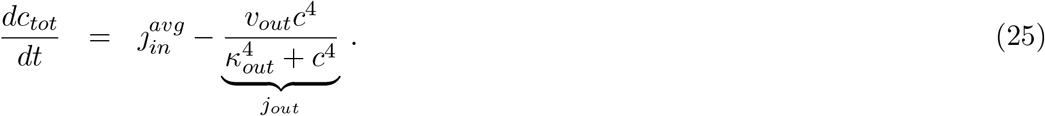

Fig. 6 (top) shows the relaxation oscillator dynamics of this open cell model using standard parameters (Table 2) and 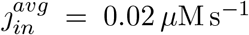. The oscillation period is on the order of minutes and the system spends most of its time in the *down* state with low cytosolic Ca^2+^ (0.05–0.11 *µ*M) and slowly increasing total cell [Ca^2+^], *c*_*tot*_, which implies slowly increasing ER [Ca^2+^]. The phase plane of Fig. 6 (bottom left) shows the *c* and *c*_*tot*_ nullclines that are found by setting the left sides of Eqs. 24–25 to zero. The *c* nullcline is *N* -shaped and has two extrema (knees). The *c*_*tot*_ nullcline is a vertical line located at the value of *c* which solves 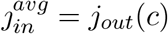. Note how the separation of time scales for *c* and *c*_*tot*_ leads to a periodic solution (black trajectory) that tracks the lower or upper branch of the *c* nullcline except for two brief excursions between branches when the trajectory passes over a knee of the *c* nullcline. Fig. 6 (bottom right) shows a bifurcation diagram for Eqs. 24–25. Type-2 relaxation oscillations that originate via Hopf bifurcations are observed for a range of average Ca^2+^ influx rates 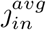. Given the separation of time scales, oscillations occur for most values of 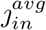 that cause the *c* nullcline to intersect the *c*_*tot*_ nullcline between the knees.

**Figure 6:**
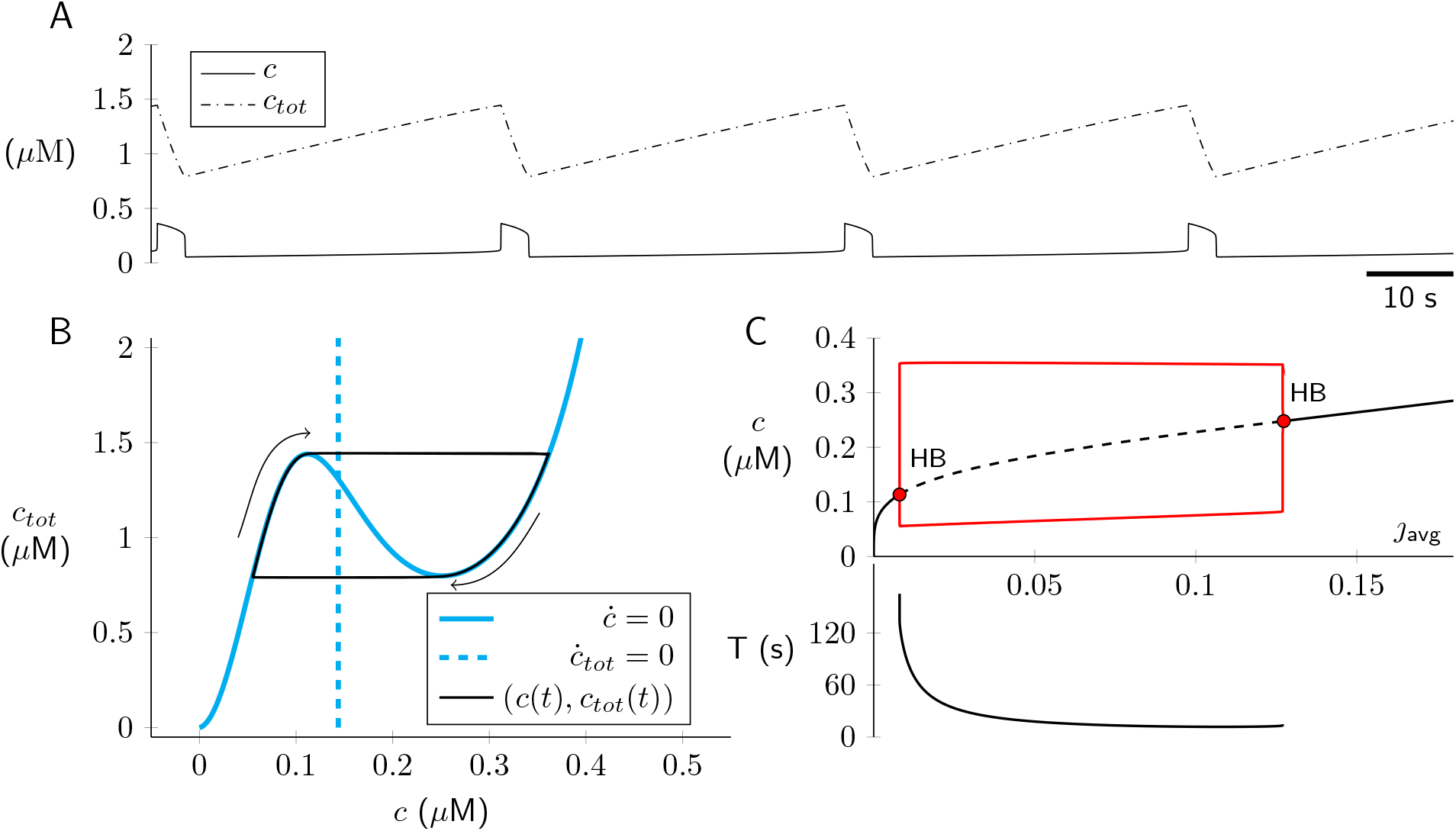
Open cell Ca^2+^ dynamics given by solution of Eqs. 24–25. Parameters: 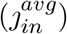 *µ*M s^*−*1^ and as in Table 2. (A) Type-2 relaxation oscillations (*c* fast, *c*_*tot*_ slow). (B) Phase plane, with *c* and *c*_*tot*_ nullcines (cyan solid and broken lines, respectively) and periodic solution (black). (C) Bifurcation diagram with 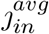 (average Ca^2+^ influx rate) as bifurcation parameter. *T* is period of relaxation oscillation.

## 5 Coupling of the inspiratory and sigh rhythms

In the model of inspiratory and sigh rhythmogenesis (Eqs. 1–11), the fast (*a, s, θ*) and slow (*c, c*_*tot*_) subsystems are bi-directionally coupled in a manner that creates the inspiratory/sigh dynamics (Fig. 1A). Fast to slow: Episodic network activity (*a*) influences dynamics of intracellular Ca^2+^ via the plasma membrane influx rate *j*_*in*_ = *J*_0_ + *J*_*a*_*a* (Eqs. 4–5). Slow to fast: Cytosolic Ca^2+^ activates a cationic current (Picardo et al., 2019) whose influence is modeled abstractly using a two-term network activity function,

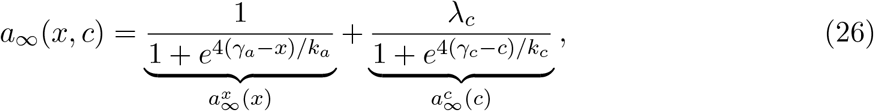

where 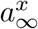 and 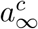 depend on synaptic drive (*x*) and cytosolic Ca^2+^ (*c*), respectively. Thus, in the model of inspiratory and sigh rhythmogenesis, the steady-state network activity function in Eq. 1 is

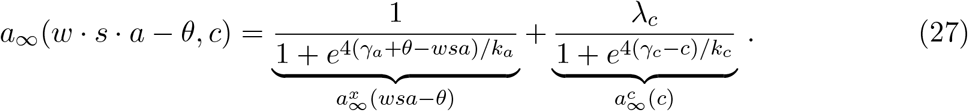

Fig. 7 plots the network activity nullcline 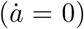,

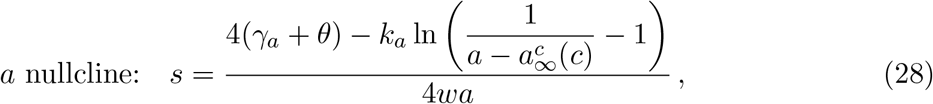

for a range of cytosolic [Ca^2+^] (0.05 *≤ c ≤* 0.35 *µ*M). Upon ER Ca^2+^ release and during the active phase of the Ca^2+^ oscillation, an upward shift of the *a* nullcline (magenta) accounts for the increase in network activity mediated by Ca^2+^-activated cationic current (Fig. 7).

**Figure 7:**
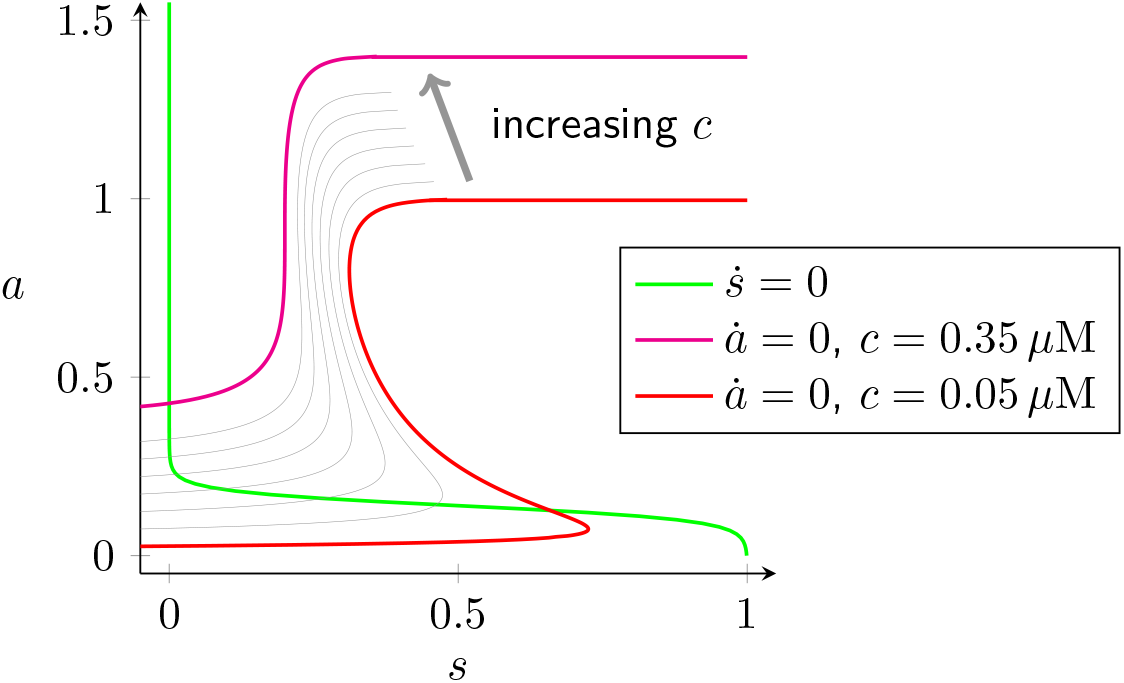
The influence of cytosolic [Ca^2+^] (*c*, gray arrow) on the network activity nullcline (*å* = 0) given by *a* = *a*_∞_(*wsa − *θ*, c*) (Eq. 27). For fixed *θ* and *c* (legend), the nullcline *a* = *a*_∞_(*s*) is given by Eq. 28. Parameters: *w* = 1, *θ* = 0.18 *γ*_*a*_ = 0.48, *λ*_*c*_ = 0.8, *γ*_*c*_ = 0.35, *k*_*c*_ = 0.05 and as in Tables 1–2. This gives 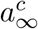 for *c* = 0.05 to 0.35 *µ*M.

## 6 Spiking model of inspiratory rhythm

Sections 2–5 described the network activity model of inspiratory rhythmogenic preBötzinger complex neurons and the dependence of sigh breathing rhythm on intracellular Ca^2+^ oscillations. The activity model approach can be compared and contrasted with models that include spiking and Ca^2+^ dynamics for *N* distinct neurons with excitatory interactions mediated by a physiologically plausible network topology. In Fig. 13, the voltage activity of each neuron is represented using the *theta model*, i.e., the Ermentrout-Kopell canonical model (Ermentrout and Terman, 2010; Ermentrout and Kopell, 1986),

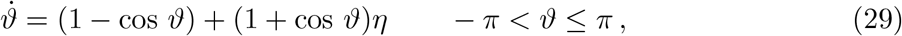

where the state variable *ϑ* is an angle in radians (not to be confused with usage of *θ* for cellular adaptation in Sections 2–5). The state space is circular (i.e., *−π* and *π* are identified) and the model produces a spike when *ϑ* increases across *π*. The parameter *η* determines the intrinsic behavior of the model neuron (see Fig. 8). For *η <* 0 the neuron is excitable; there is a stable fixed point (*ϑ*^*−*^ < 0, rest state) and unstable fixed point (*ϑ*^+^ > 0, threshold for action potential) where *ϑ*^*±*^ = *±* cos^*−*1^((1 + *η*)*/*(1 *− η*)). For *η >* 0 the neuron spikes repetitively with period 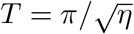 and frequency *f* = 1/*T*.

**Figure 8:**
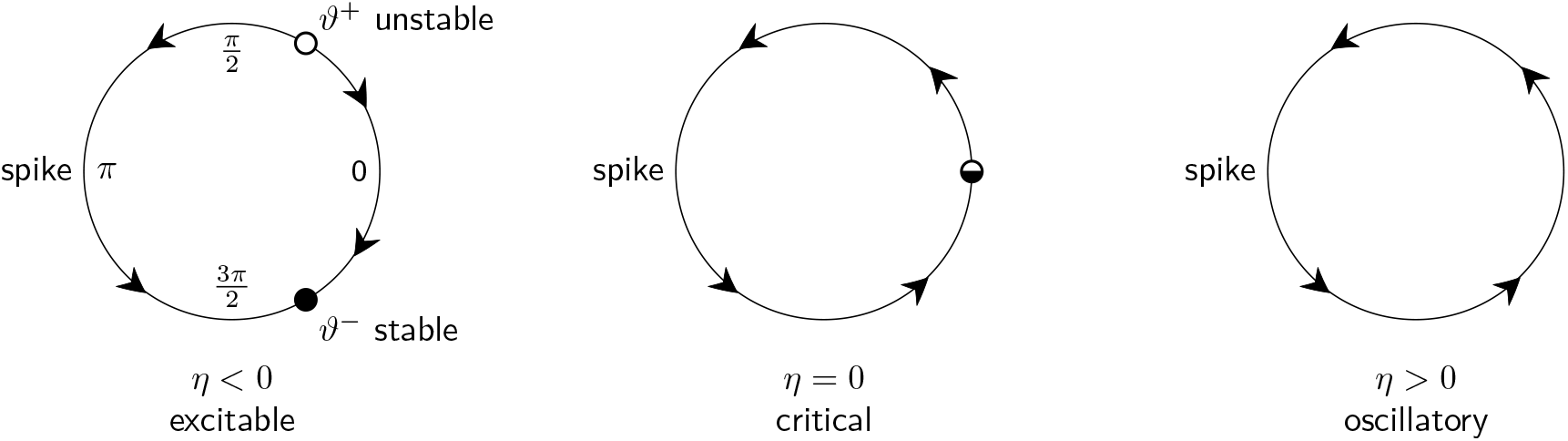
The Ermentrout-Kopell canonical model (Eq. 29) is excitable, critical, or oscillatory depending on the sign of the bifurcation parameter *η*.

Beginning with Eq. 29, a network model of inspiratory rhythmogenesis composed of *N* spiking neurons may be constructed as follows. The oscillatory phase of the *i*th neuron is given by

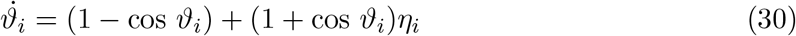

where 1 ≤ *i* ≤ *N*. The excitatory drive for the *i*th neuron takes the form

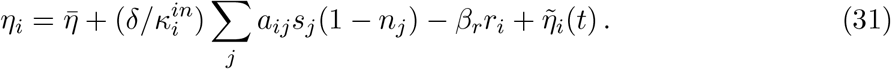

The gating variables in the above expression solve

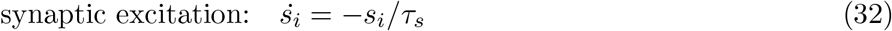

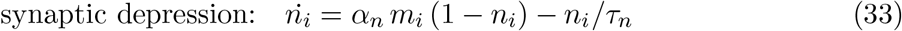

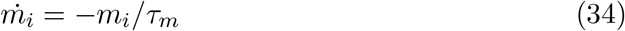

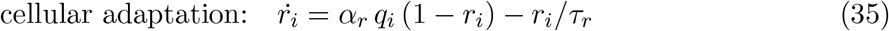

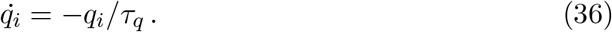

In these equations, the excitatory drive for each neuron (*η*_*i*_) is given by a sum over presynaptic neurons (Eq. 30). The state variables *s*_*i*_, *n*_*i*_, *m*_*i*_, *r*_*i*_ and *q*_*i*_, which take values between 0 and 1. The factor *s*_*j*_(1 − *n*_*j*_) in Eq. 31 is the synaptic activity of the *j*th neuron (*n*_*j*_ = 0 represents the absence of depression). The amount of cellular adaptation is given by *β*_*r*_*r*_*i*_. When the *i*th neuron produces a spike, the values of *s*_*i*_ and *m*_*i*_ are both incremented as follows:

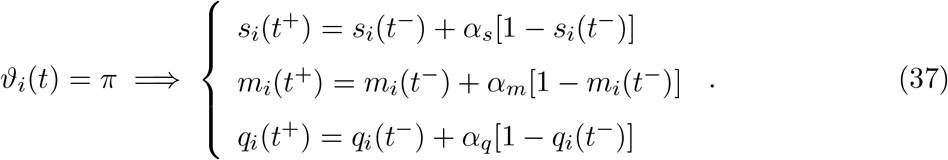

Note that *n*_*i*_ relaxes to 1 (complete depression) with exponential rate constant *α*_*n*_*m*_*i*_ so that synaptic depression is not instantaneous. In a similar manner, the cellular adaptation variable *r*_*i*_ relaxes to 1 with rate constant *α*_*r*_*q*_*i*_.

In Eq. 30, the network connectivity is given by an *N* × *N* adjacency matrix *A* = (*a*_*ij*_). The element *a*_*ij*_ > 0 if neuron *i* is postsynaptic to neuron *j* and zero otherwise. Neurons are not self-excitatory (*a*_*ii*_ = 0). The in- and out-degrees of the *i*th neuron are given by 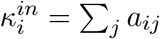 and 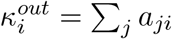, respectively. For a given simulation, the network connectivity is modeled as random directed Erdös-Réyni graph (Bollobás, 2001) with standard parameters *N* = 400 and *p* = 0.065 (see Table 3). This value for *p* greatly exceeds the threshold for connectedness given by (ln *N*)*/N ≈* 0.015. The expected value for the in- and out-degree of each neuron is *κ* = *p*(*N −* 1) *≈* 24.

**Table 3.** Parameters for the spiking network model of inspiratory rhythmogenesis.

| Symbol | Definition | Value | Units |
| --- | --- | --- | --- |
| $N$ | network size (number of neurons) | 400 | - |
| $p$ | probability that neuron $i$ excites neuron $j$ ($i \neq j$) | 0.065 | - |
| $\delta$ | strength of recurrent excitation | 0.09 | - |
| $\bar{\eta}$ | mean excitatory drive (negative for intrinsic excitability) | -0.0019 | - |
| $\nu$ | variance of stochastic excitatory drive | 0.0049 | - |
| $\alpha_s$ | growth fraction for $s$ | 1 | $\text{ms}^{-1}$ |
| $\tau_s$ | time constant for decay of $s$ | 10 | ms |
| $\alpha_m$ | growth fraction for $m$ | 0.4 | $\text{ms}^{-1}$ |
| $\tau_m$ | time constant for decay of $m$ | 300 | ms |
| $\alpha_n$ | growth fraction for $n$ | 0.011 | $\text{ms}^{-1}$ |
| $\tau_n$ | time constant for decay of $n$ | 1300 | ms |
| $j_0$ | constant $\text{Ca}^{2+}$ influx rate | 0.05 | $\mu\text{M s}^{-1}$ |
| $j_\eta$ | $\text{Ca}^{2+}$ influx rate proportionality constant | 5 | $\mu\text{M s}^{-1}$ |
| $\alpha_c$ | event-triggered $\text{Ca}^{2+}$ influx | 0.01 | nM |
| $v_{out}$ | maximum $\text{Ca}^{2+}$ efflux rate | 1 | $\mu\text{M s}^{-1}$ |
| $\lambda_c$ | maximum $\text{Ca}^{2+}$-dependent increase in synaptic drive | 0.01 | - |
| $\gamma_c$ | threshold for $\text{Ca}^{2+}$-dependent increase in synaptic drive | 0.2 | $\mu\text{M}$ |
| $k_c$ | reciprocal slope of $\text{Ca}^{2+}$-dependent increase in synaptic drive | 0.02 | $\mu\text{M}$ |

The excitatory drive for the *i*th neuron, 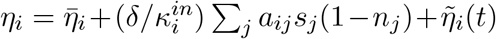, has three terms. The first term, 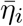 is chosen negative so that all neurons intrinsically excitable (see Fig. 8). To introduce neuronal heterogeneity, the 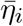 are chosen as normal random variates with mean 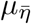 and variance 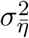. In the second term, the parameter *δ* sets the magnitude of recurrent excitation in the network, and the factor 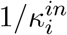 scales the synaptic interactions so that every neuron receives the same amount of excitation when all of its presynaptic neurons are active. This ensures that the network dynamics will not depend on network size provided *N* is sufficiently large. The third term is Gaussian white noise with mean zero 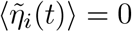 and two-time covariance 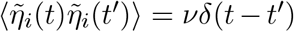. Each neuron receives an independent stream of additive noise, that is, 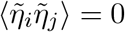 for *i* ≠ *j*.

Fig. 9 shows a raster plot and network activity for a representative simulation of the spiking model of inspiratory rhythmogenesis (Eqs. 29–37). Fig. 10 shows the bursts of spikes of patch-clamped inspiratory interneurons alongside the activity of individual model neurons in the rhythmic network simulation. The spiking network model shows good agreement with *in vitro* recordings (compare Fig. 10B to A).

**Figure 9:**
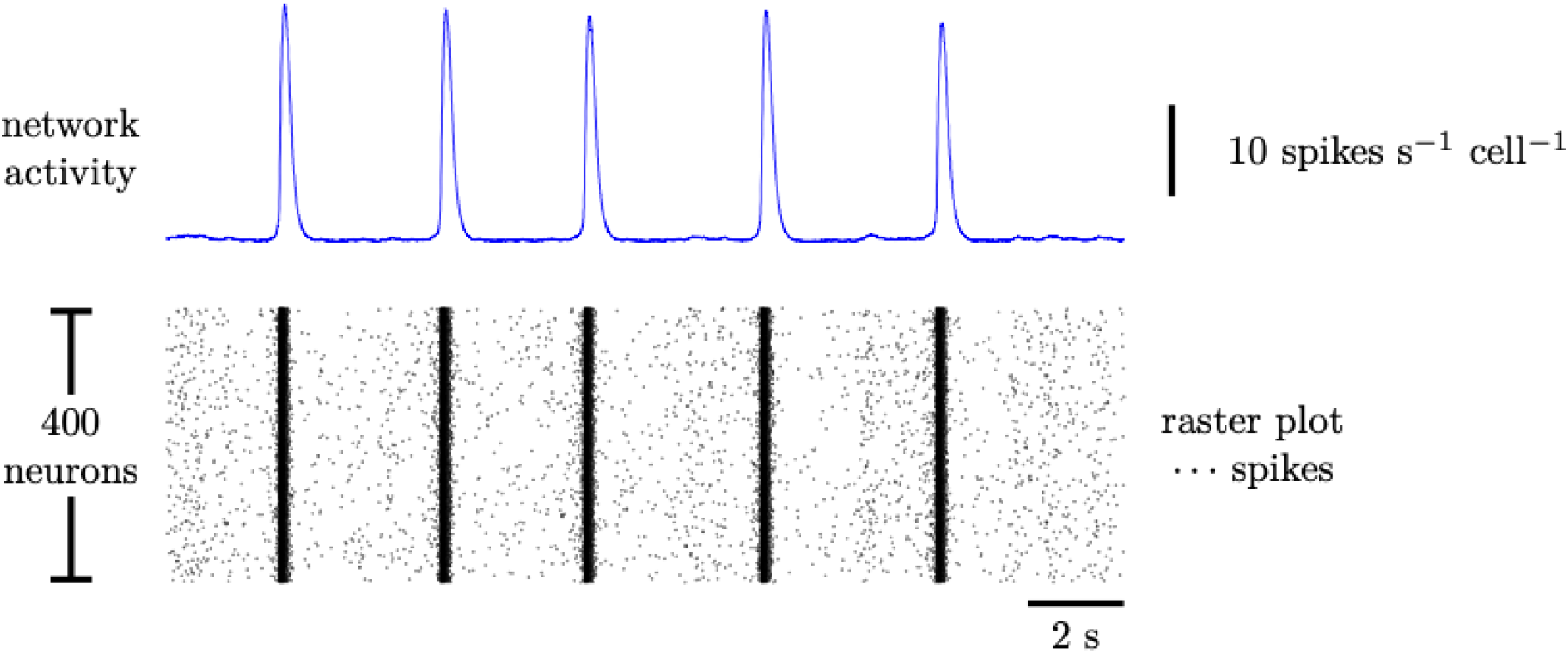
The spiking model of inspiratory rhythmogenesis (Eqs. 29–37) with Erdös-Rényi-type network connectivity with *N* = 400 and *p* = 0.065 (see Section 6). Parameters as in first block of Table 3. Cellular adaptation is not included (Eqs. 35–36 are unused and *β*_*r*_ = 0 in Eq. 31).

**Figure 10:**
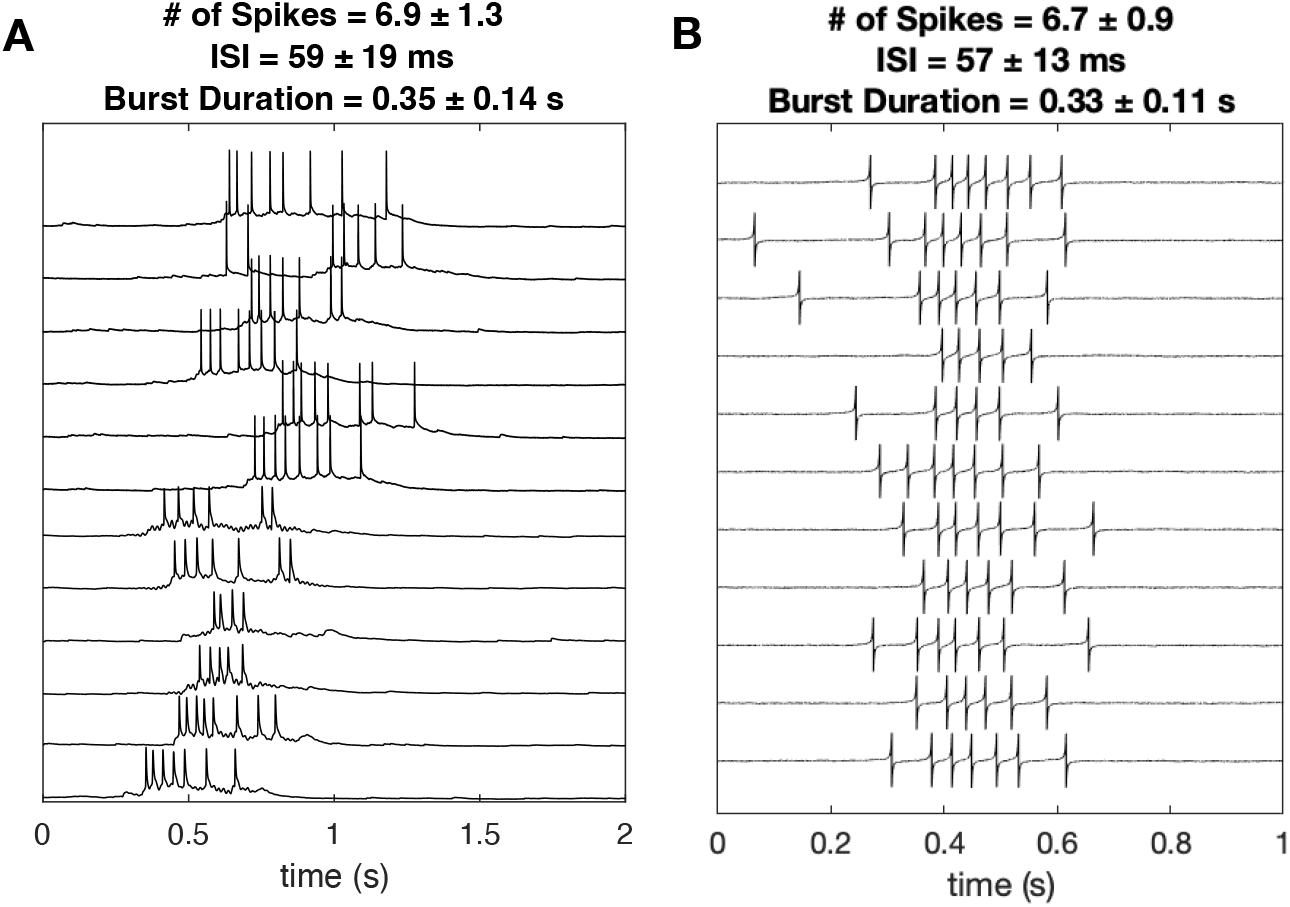
Activity of an preBötC neurons. (A) *In vitro* recordings courtesy Prajkta S. Kallurkar. (B) *In silico* spiking model simulation. Parameters as in Fig. 9.

In conditions of low excitability, the spiking network model exhibits burstlets (Fig. 11A, orange triangle) as well as bursts. Increased network excitability leads to increased burst frequency and a decrease in burstlet fraction (compare Fig. 11A and B). Both of these simulation results are consistent with experiment (Kallurkar et al., 2020; Kam et al., 2013b).

**Figure 11:**
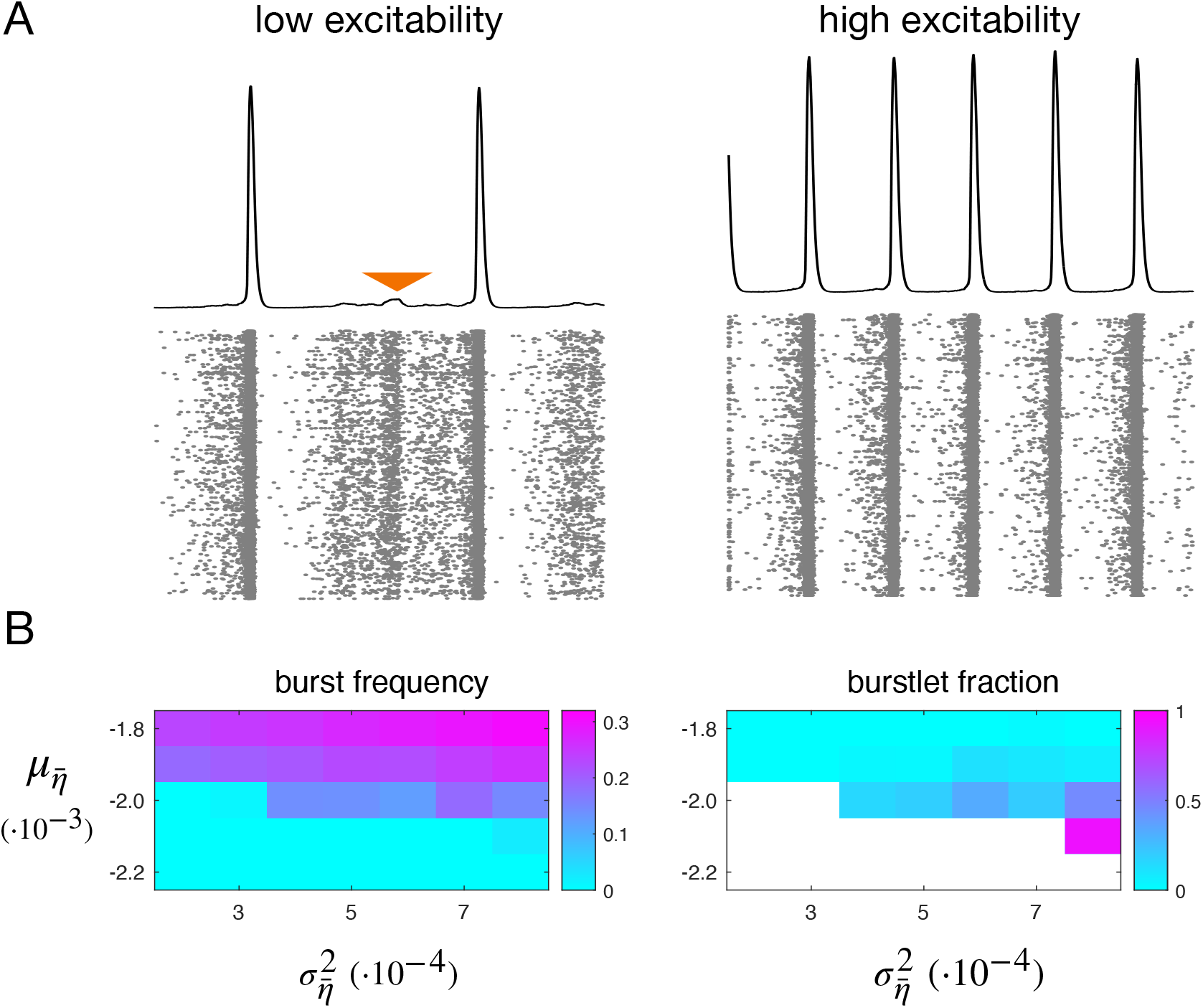
Network frequency and burstlet fraction. (A) Network rhythm at low and high excitability. (B) Burst frequency (Hz) and burstlet fraction as functions of mean 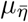 and variance 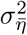 of neuronal excitability. Cellular adaptation parameters: *β*_*r*_ = 0.005, *α*_*q*_ = 0.1 ms^*−*1^, *τ*_*q*_ = 800 ms, *α*_*r*_ = 0.004 ms^*−*1^, *τ*_*r*_ = 1000 ms). Other parameters: *α*_*m*_ = 0.3 ms^*−*1^, *τ*_*n*_ = 800 ms, *J*_0_ = 0.002 *µ*M s^*−*1^ and as in Table 3.

The spiking network model is also able to reproduce glutamate uncaging experiments wherein stimulation of less than a dozen preBötC neurons elicited an ectopic network burst (Kam et al., 2013a). Fig. 12 shows raster plots of representative simulated glutamate uncaging that evokes bursts in seven neurons (both successful and failed stimulation events are shown).

**Figure 12:**
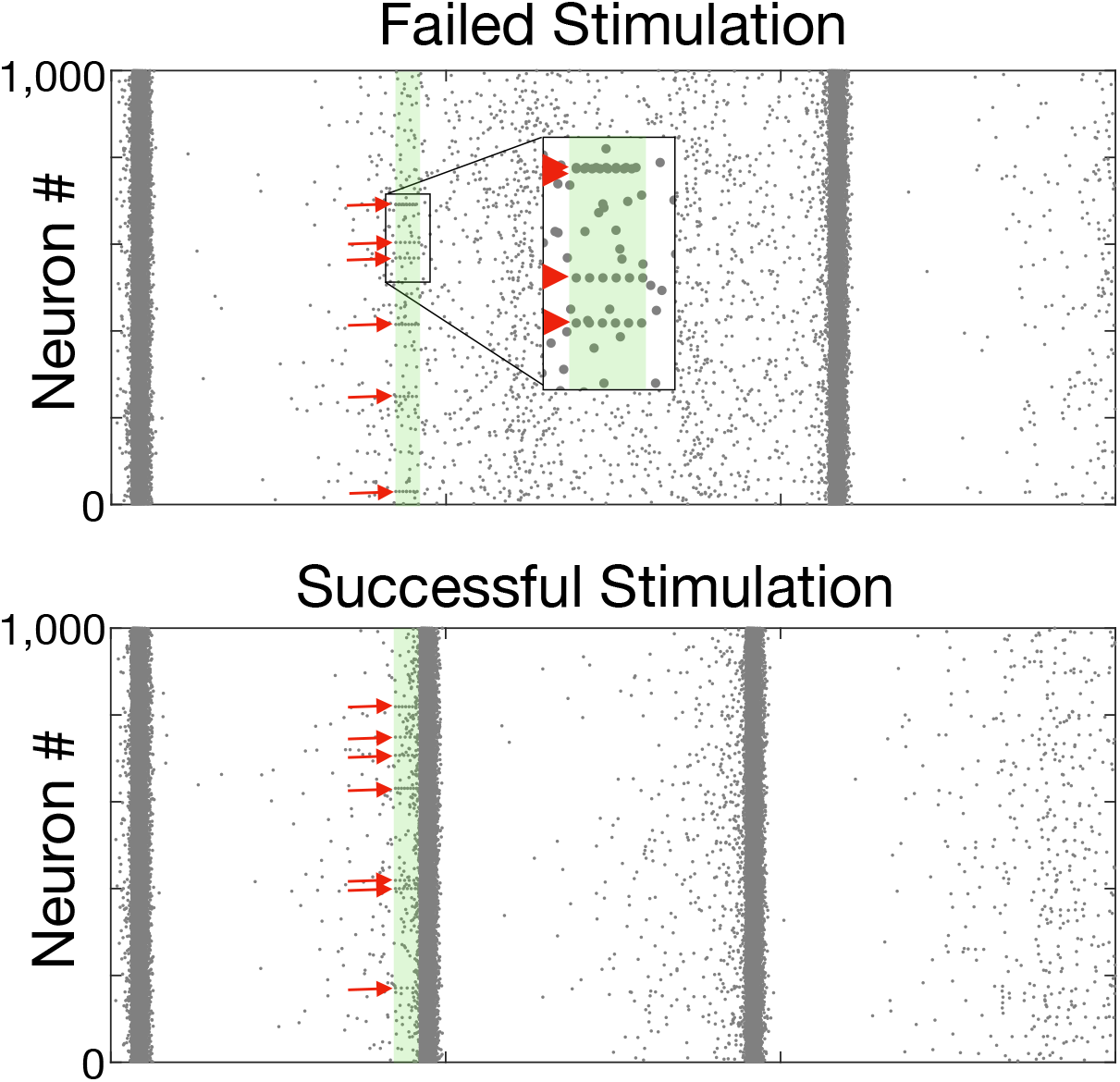
In simulated glutamate uncaging experiments, stimulation of preBötC interneurons may trigger—or fail to trigger—network bursts. Red arrows indicate seven randomly selected neurons that are driven to burst by direct stimulation.

**Figure 13:**
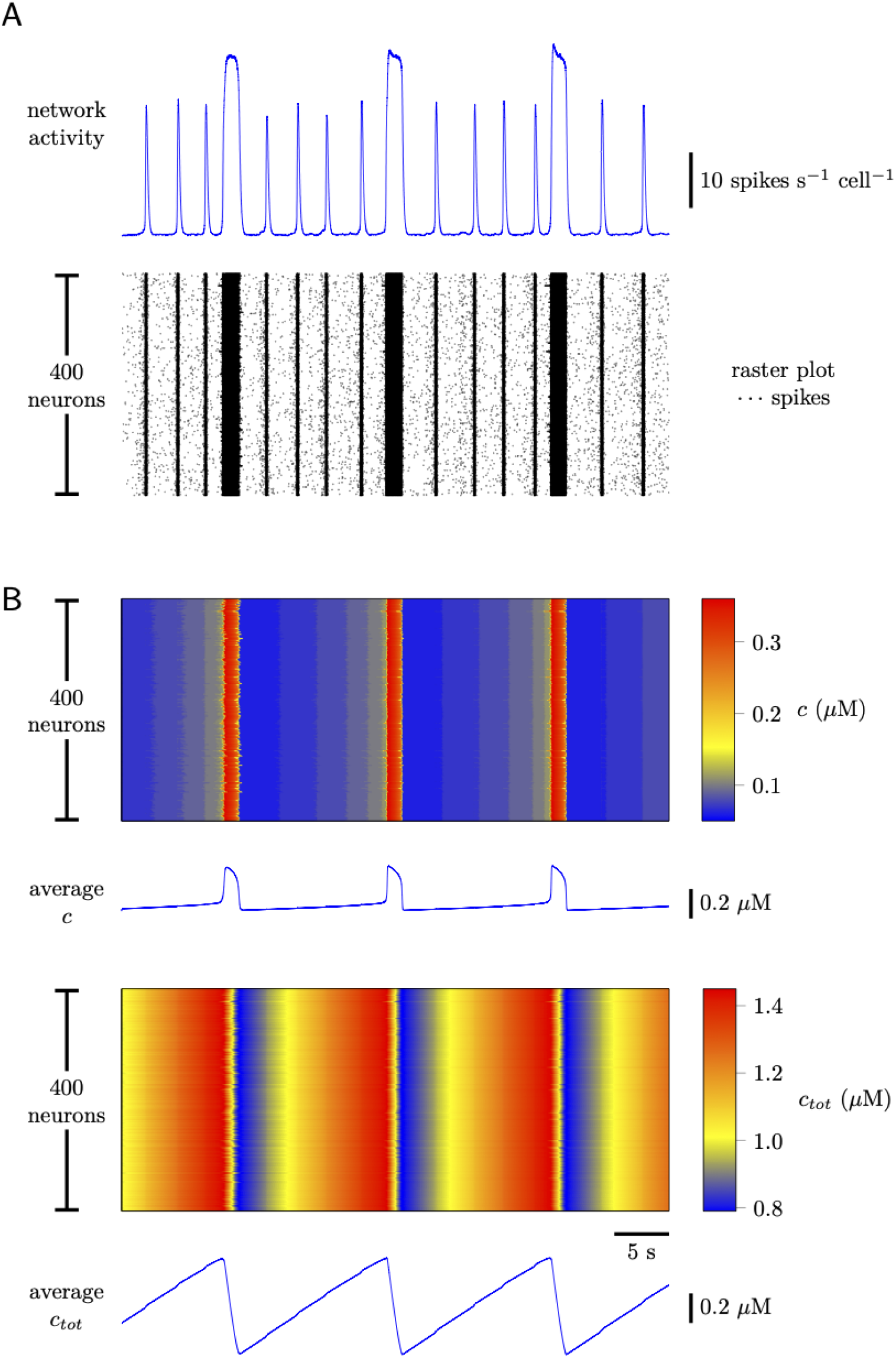
Network activity and dynamics of intracellular [Ca^2+^] for a spiking model of eupnea and sigh rhythmogenesis. The network structure is Erdös-Rényi-type with 400 neurons and 6.5% probability that any given neuron is postsynaptic to any other. (A) Network activity and raster plot. (B) Cytosolic [Ca^2+^] (*c*) and total [Ca^2+^] (*c*_*tot*_) are highly synchronized across the 400 neurons. Parameters as in top block of Table 2 and top and bottom blocks of Table 3.

## 7 Spiking network model with Ca^2+^ oscillations and sigh rhythm

To include the sigh rhythm in the spiking network model, Eqs. 30–37 are augmented with intracellular Ca^2+^ dynamics for each neuron,

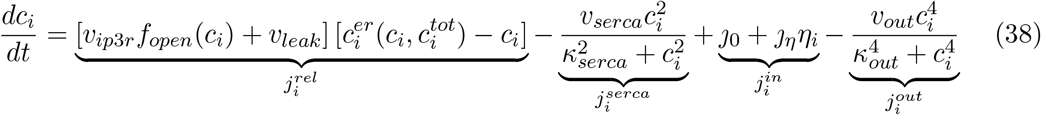

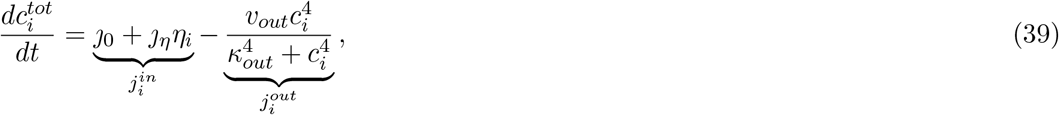

where 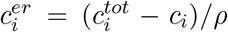. In these equations, *c*_*i*_ and 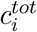 denote the cytosolic and total [Ca^2+^] of the *i*th neuron (compare Eqs. 20–21). The Ca^2+^ influx term, 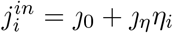, that appears in Eqs. 38 and 39 is a linear function of the synaptic activity of those neurons that are presynaptic to the *i*th neuron (*η*_*i*_). In addition, the cytosolic [Ca^2+^] of the *i*th neuron is incremented when the *i*th neuron spikes,

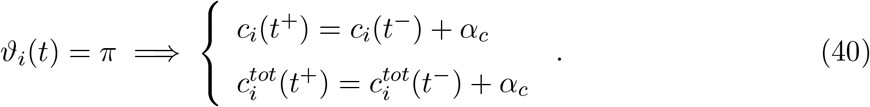

The excitatory drive for the *i*th neuron depends on cytosolic [Ca^2+^] as follows (cf. Eqs. 26 and 30),

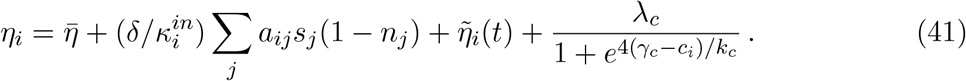

Fig. 13 shows network activity and dynamics of intracellular [Ca^2+^] for the spiking model of eupnea and sigh rhythmogenesis (Eqs. 30–41) with parameters as in Tables 2 and 3. The initial values of *c*_*tot*_ were normally distributed with mean 1 *µ*M and standard deviation 0.01 *µ*M to illustrate that synaptic activity-dependent calcium influx (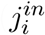 in Eq. 39) can maintain synchronous Ca^2+^ oscillations. Fig. 13 may be compared to Fig. 14, which uses identical parameters save *J*_*η*_ = 0. Comparing panel B of each figure (in particular, the third Ca^2+^ release event) shows how synaptic activity-dependent Ca^2+^ influx (*J*_*η*_ > 0) promotes Ca^2+^ subsystem synchronization among the 400 neuron population.

**Figure 14:**
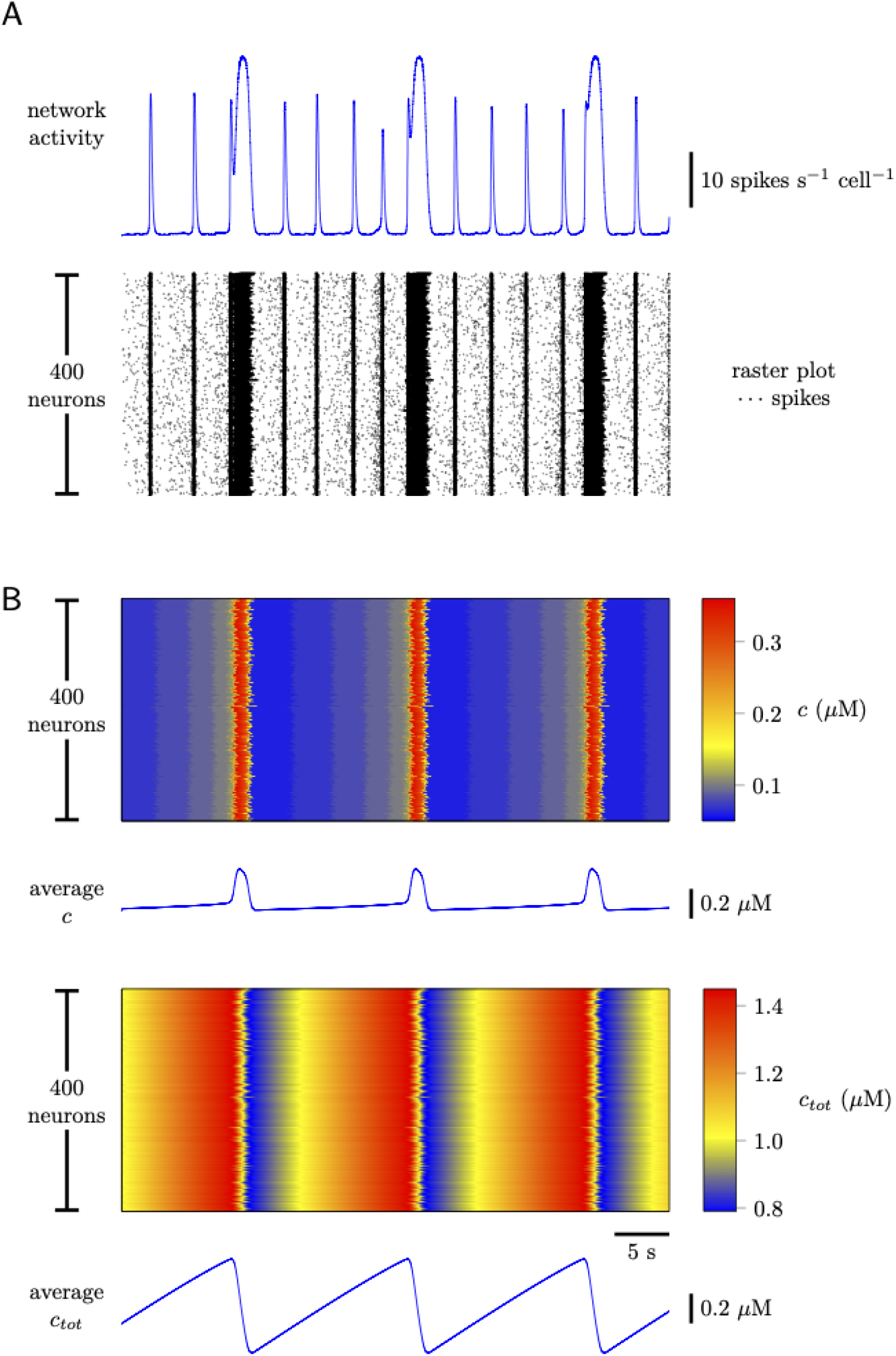
Network activity and dynamics of intracellular [Ca^2+^] for a spiking model of eupnea and sigh rhythmogenesis. The network structure is Erdös-Rényi-type with 400 neurons and 6.5% probability that any given neuron is postsynaptic to any other. (A) Network activity and raster plot. (B) In the absence of synaptic activity-dependent Ca^2+^ influx, cytosolic [Ca^2+^] (*c*) and total [Ca^2+^] (*c*_*tot*_) remain slightly desynchronized across the 400 neurons (cf. Fig. 13). Parameters: as in Fig. 13 with *J*_*η*_ = 0.

## 8 Conclusion

The model development described in this report shows that the two breathing-related rhythms generated by the preBötC (inspiration and sighs) can be generated by a single neuronal population. The simulations results presented here corroborate the hypothesis that these two disparate rhythms emerge through recurrent excitation, synaptic depression, cellular adaptation, and intracellular calcium oscillations. Both the activity model and the spiking network model described here reproduce important empirical observations. This recommends them as starting points for further study of underlying mechanisms and cellular origins of inspiratory breathing rhythms.

